# Increased circulating CD39+FoxP3+CD4+ regulatory T cells in Early Rheumatoid Arthritis facilitate the antiinflammatory action of methotrexate and associate with the clinical response

**DOI:** 10.1101/2023.10.10.561686

**Authors:** Alejandro Villalba, Laura Nuño, Marta Benito-Miguel, Beatriz Nieto-Carvalhal, Marta Novella-Navarro, Irene Monjo, Diana Peiteado, Sara García-Carazo, Alejandro Balsa, María-Eugenia Miranda-Carús

**Affiliations:** Department of Rheumatology, Hospital Universitario La Paz-IdiPaz, Madrid, Spain; Fundación San Juan de Dios, Centro de Ciencias de la Salud San Rafael, Dept. of Physiology, Universidad de Nebrija, Paseo de La Habana, 70, 28036 Madrid, Spain

**Keywords:** Early Rheumatoid Arthritis, Treg cells, CD39, adenosine, methotrexate

## Abstract

**OBJECTIVES:** FoxP3+ regulatory T cells (Tregs) are key to the immune system homeostasis; their CD39+ subset (Treg39+) hydrolises adenine nucleotides released by stressed cells, rendering the antiinflammatory adenosine. Methotrexate (MTX), inhibiting AICAR transformylase (ATIC), enhances the extrusion of adenine nucleotides and hence may help Treg39+ cells control inflammation. Therefore, we examined the relation of CD39 expression on Tregs of early RA (ERA) patients with the effect of MTX.

**METHODS:** Freshly isolated lymphocytes from 72 untreated ERA patients (duration <24 weeks) and 72 healthy controls (HCs) were examined by cytometry. Treg cell potency was assessed in cocultures of CD4+CD25+CD127- Treg with CD4+CD25- CD127+ responder T cells (Tresp).

**RESULTS:** ERA patients demonstrated a superior frequency of circulating Tregs containing increased proportions of Treg39+ cells. Total ERA Tregs were more potent than HC Tregs and MTX further heightened their potency, with greater amplification in ERA vs HC; differences were reduced by adenosine pathway blockade. The potency of isolated Treg39+ and its enhancement by MTX were comparable for ERA and HC suggesting that the differences seen with total Tregs are due to the increased ERA Treg39+ frequency. Basal Treg39+ cell proportions > 39.3 associated with a good 12 month EULAR response [RR 13.4 (2.9-75.6)]. At 12 months, the ERA Treg39+ frequency had decreased to HC levels but its association with the clinical response remained.

**CONCLUSION:** MTX cooperates with Treg39+ cells and the basal Treg39+ frequency is a predictor of clinical response. The increased circulating Treg39+ cells in untreated ERA would further facilitate the action of MTX thereby providing a slot for prompt MTX initiation.

## BACKGROUND

FoxP3+ regulatory CD4+ T cells (Tregs) are essential for maintaining the homeostasis of the immune system [1,2] and alterations in their number or function have been associated with various autoimmune conditions [2–5] including Rheumatoid Arthritis (RA) [2,6–12].

Stressed or damaged cells at inflammatory foci release adenine nucleotides to the extracellular space [13] where they act as enhancers of inflammation [13] but are rapidly hydrolysed by the combined action of ectonucleotidases CD39 and CD73 rendering the antiinflammatory agent adenosine (ADO) [13]. CD39, the rate-limiting enzyme in this cascade [14], is highly expressed by a subset of human FoxP3+Tregs (Treg39+) [8,14,15] and hydrolisis of proinflammatory adenine nucleotides seems to be an important mediator of the Treg39+ suppressive action [14,15].

Methotrexate (MTX) remains the cornerstone of treatment in Rheumatoid Arthritis (RA) [16]. Inhibition of AICAR transformylase (ATIC) by MTX results in an enhanced release of adenine nucleotides to the extracellular space [17,18] thereby providing an augmented pool of ADO precursors to be metabolized by Treg39+ cells; hence MTX may act synergistically with Treg39+ cells in the control of inflammation [8,17]. Studies in animal models of inflammation indicate that Tregs can indeed facilitate the action of MTX [8,14,15]. Furthermore, in patients with RA, the proportion of circulating Treg39+ cells has been associated with the clinical response to MTX [8,19] but there is scant information on this matter in patients with early untreated RA (ERA) [20]. In addition, there is controversy regarding the capacity of Treg39+ cells to suppress IL-17 secretion [5,21–23].

Therefore our objective was to study the frequency and function of circulating Treg39+ cells in ERA patients and its relation with the in vitro and in vivo effect of MTX.

## PATIENTS AND METHODS

### Ethics statement

The study was approved by the Hospital La Paz - IdiPAZ Ethics Committee, and all subjects provided written informed consent according to the Declaration of Helsinki.

### Patients

Peripheral blood was obtained from 72 ERA patients (Table 1) and 72 age and gender-matched healthy controls; ERA patients fulfilled 2010 American College of Rheumatology (ACR) revised criteria [24], had never received disease-modifying drugs or corticosteroids, and had a disease duration of less than 6 months (Table 1). For comparison, blood was drawn from 44 patients with longstanding RA (2010 ACR criteria) (LRA, disease duration > 2 years) (Table 1), and 44 age- and gender-matched healthy controls (Table 1). LRA patients were receiving low-dose oral methotrexate and were naïve for biological agents.

**Table 1.** Clinical data of RA patients and demographics of HC subjects.

|  | ERA<br>(n=72) | LRA<br>(n=44) | HC<br>(n=116) |
| --- | --- | --- | --- |
| Age (yrs), median (IQR) | 54 (41-60) | 61 (54-65) | 57 (43-62) |
| Female, n (%) | 58 (81) | 35 (80) | 93 (80) |
| Disease duration, median (IQR) | 16 wks (8-24) | 10 yrs (9-21) | N/A |
| Sero+ (RF and/or ACPA +), n (%) | 56 (78) | 44 (100) | N/A |
| RF+, n (%) | 46 (64) | 41 (92) | N/A |
| ACPA+, n (%) | 47 (65) | 40 (90) | N/A |
| DAS28, median (IQR) | 5.1 (4.2-5.6) | 3.8 (3.6-5.8) | N/A |
| Methotrexate | 0 | 44 (100) | N/A |
| Prednisone ≤ 5 mg/day | 0 | 6 (14) | N/A |

In addition, synovial fluid (SF) was obtained from the knee joints of 8 patients with LRA. ERA patients very rarely present with joint effusions requiring drainage and therefore no ERA SF was available for this study.

## EXPERIMENTAL METHODS

### Isolation of CD4+ T cells and purification of CD4+CD25- and CD4+CD25+ T cell fractions

Regulatory and responder T cells were sorted out as described [6]. In brief, freshly isolated mononuclear cells were separated from human blood by density centrifugation over Ficoll-Paque Plus (GE Healthcare, Chalfont St. Giles, United Kingdom). CD4+ T cells were subsequently purified by negative selection in an Automacs (Miltenyi Biotec, Bergisch Gladbach, Germany) using the “CD4+ T Cell Isolation Kit II” from Miltenyi Biotec. Isolated CD4+ T cells were 98% pure and free of detectable CD14+ monocytes, CD56+ NK and CD19+ B cells. Subsequently, CD4+CD25-CD127+ (T responder = Tresp), CD4+CD25-CD127+CD39- (Tresp39-), CD4+CD25+CD127- (T regulatory = Treg), CD4+CD25+CD127-CD39+ (Treg39+) and CD4+CD25+CD127-CD39-(Treg39-) populations were isolated from total CD4+ T cells by FACS sorting in a BD Vantage SE (Becton Dickinson) to a purity >98%.

### Stimulation for intracellular IL-17 detection in freshly isolated T cells

CD4+T cells were cultured immediately after isolation in 24-well plates (10^6^ cells/well) containing RPMI 1640 medium (Lonza) with 10% FCS, 2 mM L-glutamine, 50 U/ml penicillin, 50 mg/ml streptomycin and 50 mM 2-mercapto-ethanol (‘‘complete RPMI medium’’). T cells were stimulated for 16 h with phorbol myristate acetate (PMA) (10 nM) and ionomycin (2 mM), in the presence of 4 mM monensin (all three from Sigma-Aldrich).

### Assessment of Treg cell function

Isolated Treg were cocultured for 6 days with Tresp cells at a 1:10 Treg:Tresp ratio; previous work in our group showed that this proportion is optimal for assessing the action of agents thay may modify the Treg suppressor function [6]. Cocultures (2x10^4^ Treg:2x10^5^ Tresp cells/well) were stimulated as described [6] in flat-bottom 96 well plates with a soluble anti-CD3 IgE subclass (T3/4.E, Sanquin, Amsterdam, The Netherlands, formerly CLB). It is known that anti-CD3 antibodies, when bound to plastic culture surfaces, facilitate crosslinking that induces T cell proliferation. In contrast, soluble anti-CD3 antibodies do not induce T cell proliferation in the absence of antigen-presenting cells or anti-CD28 abs. A remarkable exception is an IgE anti CD3 antibody that was developed at the CLB as a switch variant originating from an IgG1 anti CD3 MoAb (CLB-T3/4.E). Soluble CLB-T3/4.E is able to stimulate T cells directly, without the need for a second signal [25]. No accessory cells and no costimulatory antibodies were used in an attempt to induce a mild stimulation of T cells and to avoid the use of agents that may interfere with regulatory T cell function. T cell proliferation was assessed by ^3^H Thymidine incorporation and CFSE dilution. For Thymidine assays, eighteen hours before the termination of the cultures, the plates were pulsed with 0.5 µCi/well [3H]thymidine (GE Healthcare). The cells were harvested on paper filters and [^3^H]thymidine uptake was measured in a liquid scintillation counter. To track division by cytometry in Treg/Tresp cocultures, Tresp cells were labeled before initiating the cocultures with CFSE (Molecular Probes/Invitrogen), at a final concentration of 8 µM. For detection of soluble cytokines by ELISA, PMA (10 nM) and ionomycin (2 mM) were added to the medium for the last 6 hours before terminating the cocultures and supernatants were collected and stored at -20°C. For intracellular cytokine examination, 4 mM monensin was added one hour after the addition of PMA/ionomycin, that is, 5 hours after terminating the cocultures. Examination of intracellular cytokine expression by cytometry in Tresp was done after gating for CFSE positive fluorescence. Potency of suppression was calculated as [1- (proliferation or cytokine secretion of Treg+Tresp coculture / proliferation or cytokine secretion of Tresp cells)]. Cell viability was not decreased in the presence of Treg cells as determined by flow cytometry after staining with 10 mM JC-1 (5,59,6,69-tetrachloro-1,19,3,39- tetraethylbenzimidazol-carbo-cyanine iodide, Molecular Probes - Invitrogen, Leiden,The Netherlands) to evaluate the mitochondrial membrane potential.

### In vitro treatment with adenosine (ADO) pathway blockers and/or methotrexate (MTX)

Treg/Tresp cocultures were pretreated for 5 hours with MTX (Sigma) (100 nM) before adding anti CD3ε (Sanquin) (0.25 μg/ml) without washing away MTX. This concentration was chosen as the opitmal for the assays described herein after titrating ascending doses as previously reported [6]. In some conditions, the CD39 inhibitor ARL67156 (Sigma) (10 μM) [26], the A2A ADO receptor (ADORA2A) antagonist ZM241385 [27] (Sigma) (10 nM) or the A1 ADO receptor (ADORA1) antagonist 8- cyclopentyl-dipropylxanthine (DPCPX) (10 μM) (Sigma) were added to the medium in the absence of or together with MTX.

### Culture of human fibroblasts

Human fibroblasts were obtained as described [6]. In brief, RA synovial fibroblasts (RASFib) and osteoarthritis synovial fibroblasts (OASFib) were isolated by collagenase digestion (type I; Worthington Biochemical) of human synovial tissue obtained at arthroplasty or synovectomy, after patients had signed the informed consent form approved by the ethics committee, according to the Declaration of Helsinki. Dermal fibroblasts were obtained by collagenase digestion of normal skin from punch biopsies of five healthy volunteers. Cells were plated in 75-cm2 flasks (Corning Costar) and grown in DMEM (Lonza) supplemented with 10% FCS (Lonza), 2 mM L-glutamine, 50 U/ml penicillin, and 50 mg/ml streptomycin (Lonza). Cells were passaged at 1/2 dilution when reaching 95% confluence, by gentle trypsinization (0.05% trypsin/0.53 mM EDTA; Invitrogen). Fibroblasts were used between the third and fifth passages. At this time, they appear to be a homogeneous population of fibroblast-like cells that stain positive with anti-Thy-1 (CD90) Ab and are negative for the expression of CD1, CD3, CD19, CD14, HLA class II, CD80, and CD86, as determined by flow cytometry and fluorochrome-conjugated mAbs (BD Pharmingen).

### Coculture of Treg cells with human fibroblasts

Coculture experiments were performed as described [6], with confluent fibroblast cultures prepared 48 h before contact. All experimental conditions were performed in triplicate and variation between replicates was <5%. Fibroblasts were seeded in 96-well plates (Corning) at 1x10^4^ cells per well. Forty-eight hours later, Treg cells (10^5^ cells/well) were added and the medium was supplemented with soluble anti-CD3ε [25] (T3/4.E, Sanquin). T3/4.E is able to stimulate Tresp cells directly in soluble form [25], but not isolated Treg cells since they are anergic [1,2]; it allows the evauation of stimulated T cell responses in the presence of RASFib [6]. Cocultures were maintained for 6 days. Parallel experiments included the addition of a neutralizing anti-IL-15 MoAb (10 mg/ml) (mab247, mouse IgG1, R&D systems, Abingdon, UK). Alternatively, the binding control anti-HLA class I (clone W6/32, 10 mg/ml Sigma, St. Louis, MO), an anti-human MHC class II (HLA DR, DP, DQ; 10 mg/ml, BD biosciences, San Jose, CA) or isotype control murine IgG1 (R&D Systems), were used. These antibodies were incubated with RASFib for 30 min at 37°C; T cells were subsequently added without washing. A 0.4 µm Transwell system (Corning) was used to conduct some co-culture experiments. The system consists of two compartments: a top well, with a porous matrix (0.4 µm), and a bottom well. This setup allows co-culture of two types of cells to grow in the same medium with soluble factors exchanged through the pores, while preventing direct contact between them. RASFib were grown to confluence in the bottom well, and T cells were added either to the same well (allowing contact) or in the top well (avoiding contact). To detect intracellular IL-17 expression in cocultured Tregs, PMA (10 nM), ionomycin (2 mM) and monensin (4 mM) were added to the medium for the las 6 hours before terminating the cocultures.

### Intracellular cytokine staining, surface staining, and flow cytometry

Cells were surface stained with fluorochrome-conjugated mAbs to examine the expression of the phenotypic markers CD3 (clone UCHT1, BD Pharmingen), CD4 (RPA-T4, BD), CD127 (HIL-7R-M21, BD), CD25 (2A3, BD), and CD39 (MZ18-23C8, Milteyi). For intracellular staining, cells were subsequently permeabilized with FoxP3 Staining Buffer Set (Miltenyi Biotec) according to the manufactureŕs instructions, and incubated on ice for 30 minutes with fluorochrome-labeled mAbs directed to interferon gamma (clone B27, BD), IL-17 (clone eBio64DEC17), FoxP3 (clone 236A/E7), Helios (clone 22F6) and/or Rorγt/RORC2 (clone AFKJS-9) (all from eBioscience), or isotype control mAbs. Events were acquired in a FACSCalibur or FacsCelesta flow cytometer and analyzed using FlowJo 10.8.1 software (all three from Becton Dickinson).

### Soluble cytokine detection

ELISAs for IFN-γ, TNF-α and IL-17 were performed in cell-free supernatants using kits from BD-Pharmingen.

### Statistical analysis

Comparison between groups was by Mann-Whitney or Kruskal-Wallis test. When appropriate, Bonferroni correction for multiple comparisons was applied. Correlations were analyzed using Spearman’s rank correlation coefficients. Nominal measures were analyzed using the Fisheŕs exact test. Logistic regression was applied to verify the independent associations between immunological and clinical parameters. All analyses were performed with GraphPad Prism version 9.4.1 software (GraphPad Software, San Diego, CA, USA).

## RESULTS

### Frequency of total Treg and their Treg39+ or Treg17 cell subsets in RA

Treg cells were identified as CD4+CD25+CD127-FoxP3+ (Fig. 1A), and their frequency was expanded in ERA when compared with HC subjects (Fig. 1A). Further phenotypical characterization revealed that a proportion of Treg cells expressed CD39 (Fig. 1B), and interestingly, the percentage of CD39+ cells was increased among Treg cells (Treg39+) of ERA patients (Fig. 1C). In addition, IL-17 expression was detected by cytometry among Treg cells (Fig. 2A), and strikingly the percentage of circulating Tregs that expressed IL-17 (Treg17) was decreased in ERA (Fig 2B). In contrast, the proportions of Treg cells and their subsets in LRA subjects were not different from those observed in healthy controls (Fig. 1 A, C; Fig. 2 B). Of note, the synovial fluid of LRA patients (LRASF) contained the highest proportions of total Treg and Treg39+ cells and lower percetages of Treg17 cells (Fig.1 A, C; Fig. 2 B). Treg17 cells were mostly Helios negative and coexpressed RorC2 with FoxP3 (Fig. 2 C).

**Figure 1.**
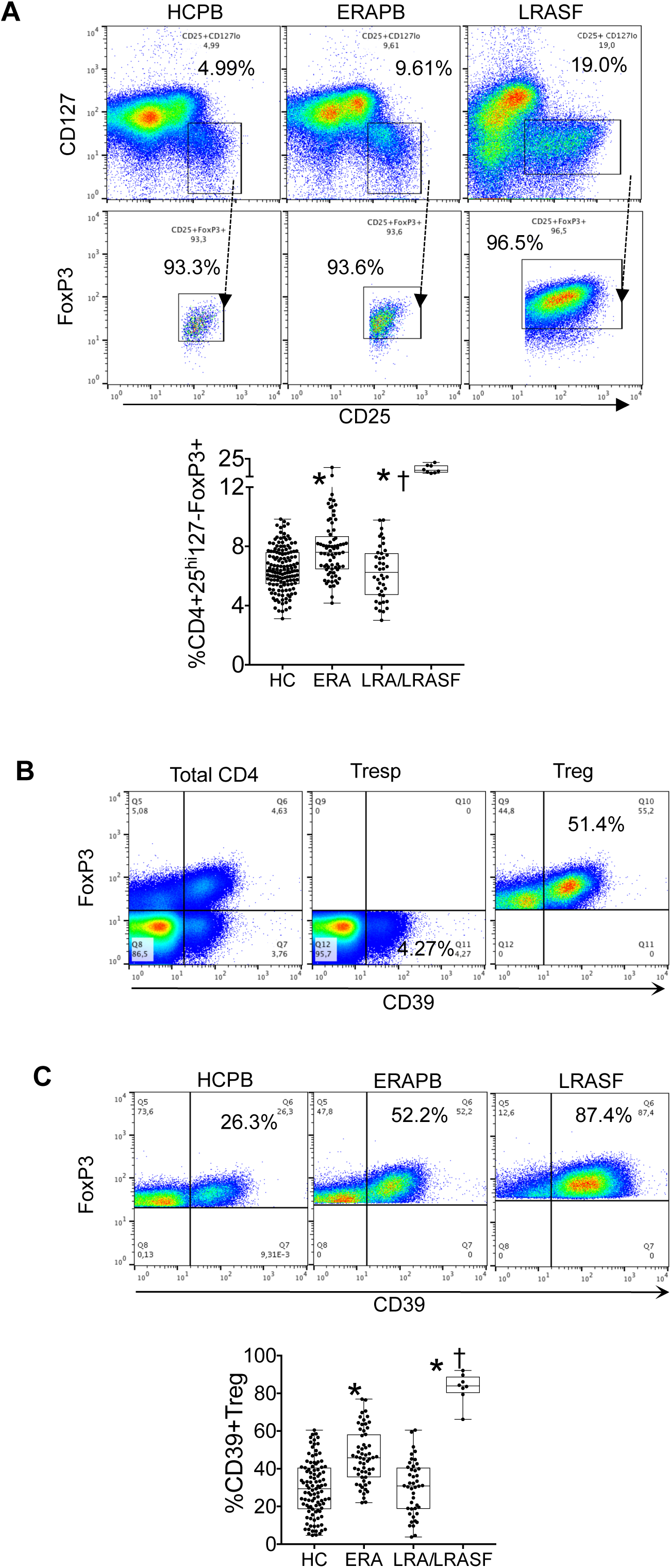
Frequency and phenotype of Treg cells in the peripheral blood and synovial fluid of Rheumatoid Arthritis (RA). **A**. Frequency of CD25+CD127-FoxP3+ CD4+ T (Treg) cells in HC (n=116), ERAPB (n=72), LRAPB (n=44) and LRASF (n=8). The Treg cell frequency was augmented in ERAPB and RASF but not in LRAPB. Representative dot plots show CD127 and CD25 expression in cells gated for CD3 and CD4 (top row), or CD25 and FoxP3 expression in cells first gated for CD3 and CD4 and then for CD25^hi^ and CD127^lo^ expression (bottom row). **B**. Representative dot plots of CD39 and FoxP3 expression in cells gated for CD3+CD4+ expression (Total CD4, left), CD3+CD4+CD25-CD127+FoxP3- expression (Tresp, middle) or CD3+CD4+CD25^hi^CD127-FoxP3+ expression (Treg, right). **C**. Increased Expression of CD39 (HC, n=116; ERAPB, n=72; LRAPB, n=44, LRASF, n=8) among Treg cells of ERAPB and RASF. CD39 was increased among Treg cells of ERAPB and RASF. Representative dot plots show CD39 or IL-17, and FoxP3 expression, in cells gated as shown above in A (first gated as CD3+CD4+CD25hiCD127- and then on CD25^hi^FoxP3+). HCPB, healthy control peripheral blood; ERAPB, early RA peripheral blood; LRAPB, longstanding RA peripheral blood; LRASF, longstanding RA synovial fluid. Graphs show individual values, with box and whiskers marking the median, interquartile range, maximum and minimum. *p<0.01 vs HC; † p<0.01 vs ERA (Mann-Whitney Test).

**Figure 2.**
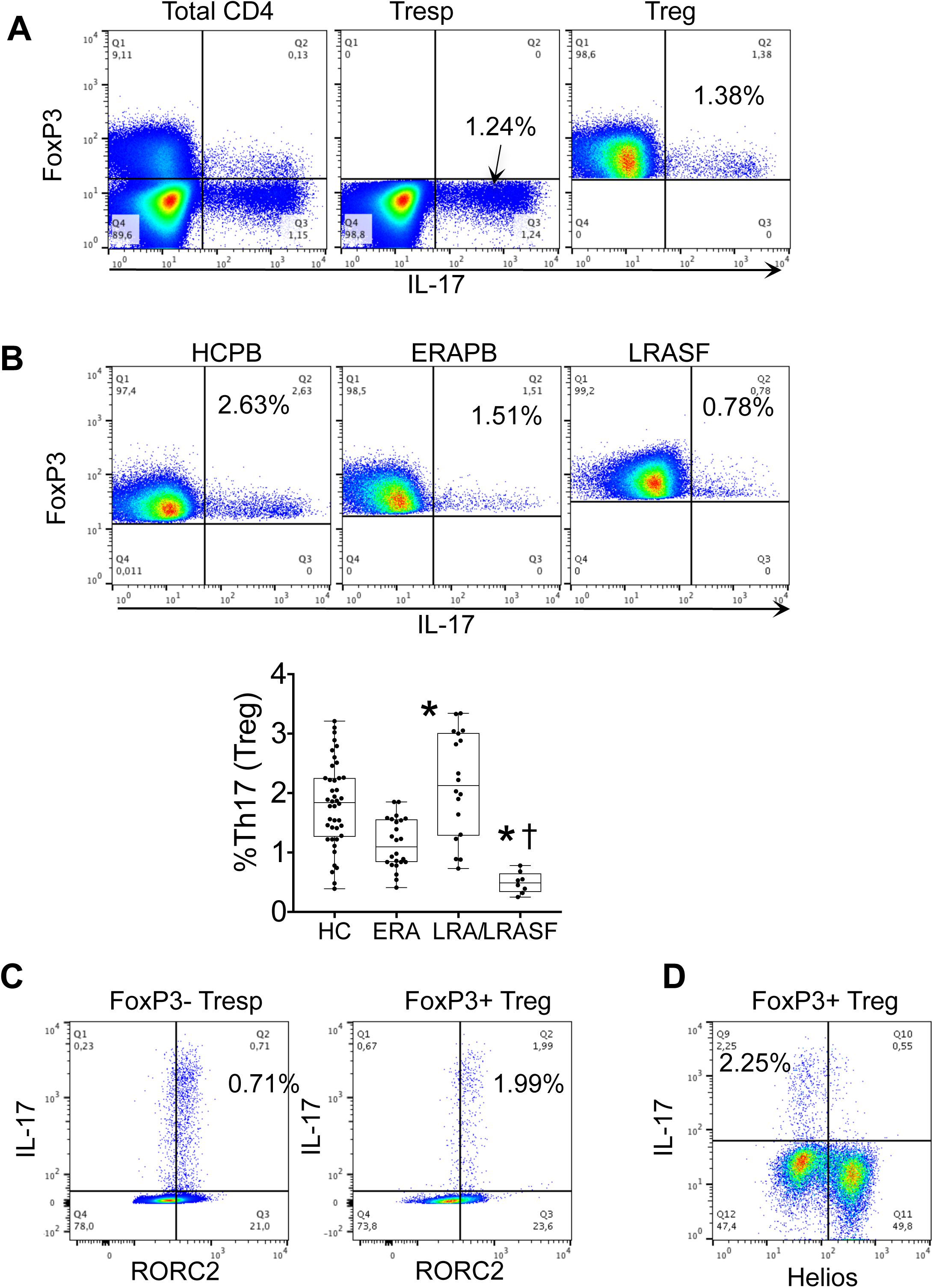
Intracellular IL-17 expression in Treg cells from the peripheral blood and the synovial fluid of Rheumatoid Arthritis (RA). **A.** Representative dot plots of IL-17 and FoxP3 expression in cells gated for CD3+CD4+ expression (Total CD4, left), CD3+CD4+CD25-CD127+FoxP3- expression (Tresp, middle) or CD3+CD4+CD25^hi^ CD127-FoxP3+ expression (Treg, right). **B**. Decreased expression of IL-17 among Treg cells of ERAPB and RASF. Representative dot plots show IL-17, and FoxP3 expression, in cells gated for CD3, CD4 and FoxP3. Graphs show individual values, with box and whiskers marking the median, interquartile range, maximum and minimum. *p<0.01 vs HC; † p<0.01 vs ERA (Mann-Whitney Test). **C.** FoxP3-Tresp and FoxP3+Treg cells that express IL-17 coexpress RORC2. Representative dot plots show RORC2 and IL-17 expression, in cells gated for CD3, CD4 and absence (left) or presence (right) of FoxP3. **D**. FoxP3+Treg cells that express IL-17 are mostly Helios negative. Representative dot plots show Helios and IL-17 expression, in cells gated for CD3, CD4 and FoxP3.mHCPB, healthy control peripheral blood; ERAPB, early RA peripheral blood; LRAPB, longstanding RA peripheral blood; LRASF, longstanding RA synovial fluid.

### Expression of IL-17 in Treg39+ and Treg39- cells

IL-17 was expressed by both Treg39+ and Treg39- cells (Fig. 3A, B), but the proportion of IL-17+ T cells was significantly lower in Treg39+ as compared with Treg39- cells (Fig. 3 A, B), and this was true for all four groups of studied samples (peripheral blood of HC, ERA or LRA and SF of LRA) (Fig. 3 A, B).

**Figure 3.**
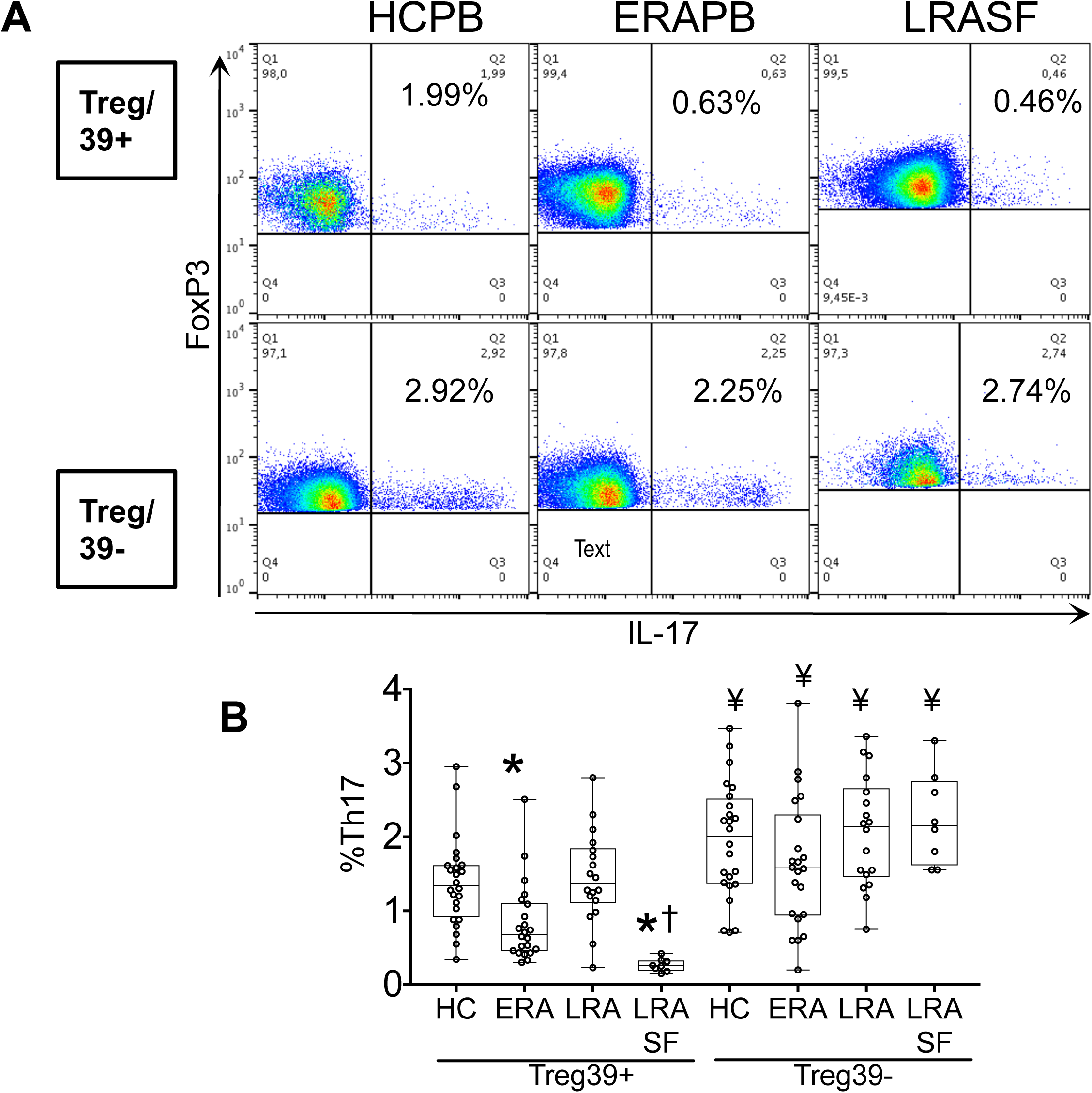
Intracellular IL-17 expression in Treg39+ and Treg39- cells from the peripheral blood and synovial fluid of Rheumatoid Arthritis (RA). **A**. Representative dot plots show IL-17 and FoxP3 expression in cells first gated as shown in figure 1 (initially as CD3+CD4+CD25^hi^CD127- and then on CD25^hi^FoxP3+) and subsequently gated for the presence of CD39 (top row) or absence of CD39 (bottom row). **B**. IL-17 expression of Treg39+ and Treg39- cells derived from the peripheral blood of HC (n=24), ERA (n=24) or LRA (n=18), and the synovial fluid of LRA patients (n=8). Graphs show individual values, with box and whiskers marking the median, interquartile range, maximum and minimum; *p<0.01 vs HC; † p<0.01 vsERA; ¥ p<0.01 vs Treg39+ cells of the same group of patients, Mann Whitney test).

In addition, as compared with HC, IL-17 expression was significantly decreased among circulating Treg39+ cells of ERA patients and maximally decreased in RASF CD39+Tregs (Fig. 3 A, B). In contrast, IL-17 expression in Treg39- was not significantly different among the four studied groups (Fig. 3 A, B). As described [6], Treg cells did not produce IFNγ or TNFα (not shown).

### Suppressive capacity of Treg cells on Tresp cell proliferation

The gold standard for Treg cell definition is demonstration of their suppressive competence. To this end, total Treg, Treg39+, Treg39- and Tresp cells were isolated from the peripheral blood of 6 HC and 6 treatment-naïve ERA patients, and from the synovial fluid of 6 LRA patients. Using a previously described system [6], we tested the Treg cell suppressive potency at a Treg/Tresp ratio of 1:10. Cell viability was not decreased in the presence of Treg cells as determined by assessment of the mitochondrial membrane potential by flow cytometry and JC-1 saining (Fig. 4A). Total Treg, isolated Treg39+ and isolated Treg39- cells were able to suppress Tresp proliferation as determined by ^3^H-Thymidine incorporation or CFSE dilution (Fig. 4 B, C), and results were comparable when using either of these methods (Fig. 4 C). As reported [6], the suppressive potency of ERA Total Treg cells was superior to that of HC Tregs (Fig.4 B, C), whereas LRASF Tregs demonstrated the highest potency (Fig.4 B, C). We herein show that Treg39+cells are significantly more potent as compared with Treg39-, and that the potency of Treg39+ or Treg39- is comparable among the groups (Fig. 4C). This suggests that the differences observed in assays using total Tregs can be attributable to the increased Treg39+ proportions present in ERA peripheral blood and LRASF.

**Figure 4.**
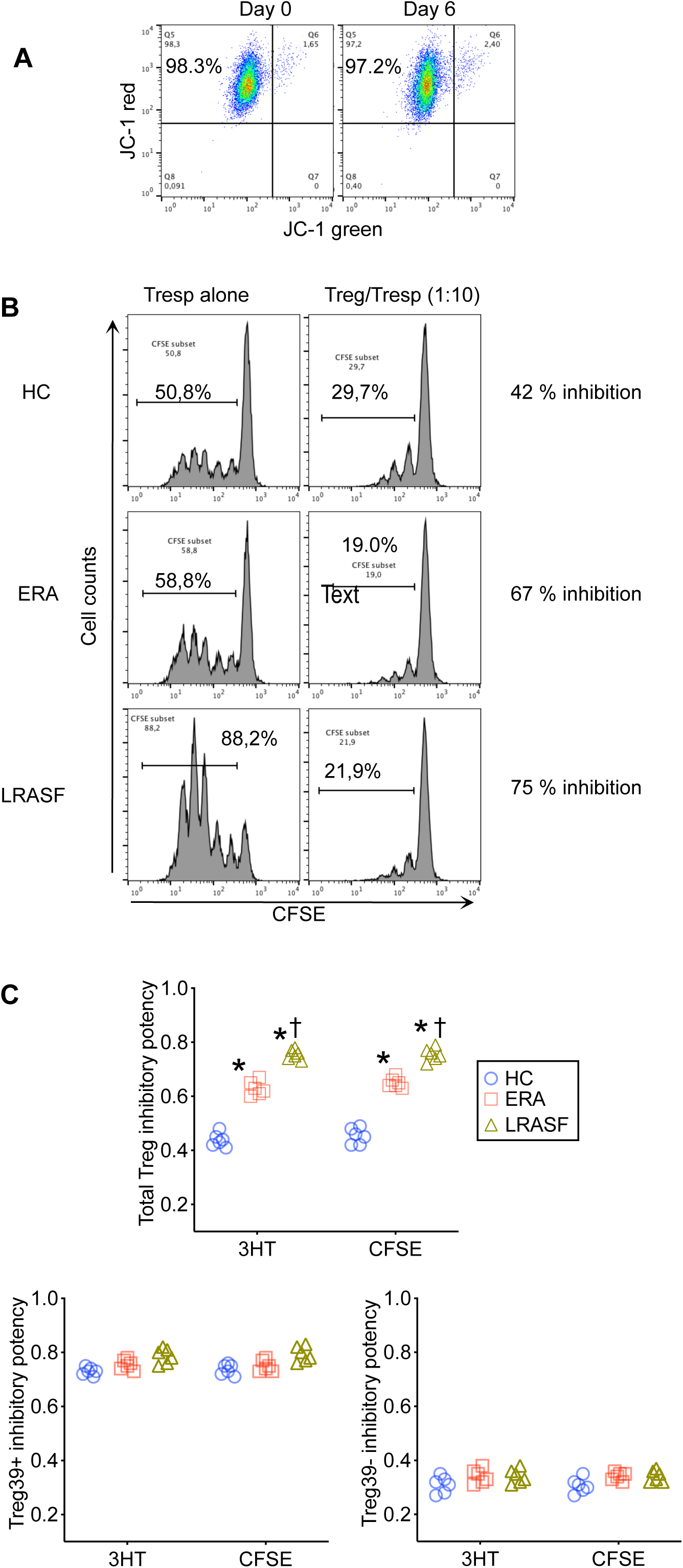
Inhibitory potency of Treg cells on Tresp cell proliferation. The potency of inhibition of Treg on Tresp cell proliferation was assessed in 6 day Treg/Tresp cell cocultures (Treg/Tresp cell ratio 1:10) using both a ^3^H Thymidine incorporation assay and CFSE dilution. **A.** Representative flow cytometry dot-plots of Treg/Tresp cell cocultures stained with JC-1 on days 0 (basal time point) and 6 (final day). Cell viability was maintained above 96% throughout. **B.** Isolated Treg were cocultured for 6 days with CFSE-labeled Tresp cells. Shown are representative flow cytometry histograms of CFSE dilution in cultures of Tresp cells alone (left column) or Treg/Tresp cocultures (1:10) (right column) **C.** Inhibitory potency on Tresp proliferation, of Total Treg, CD39+ and CD39-Treg, as assessed by ^3^H Thymidine incorporation or CFSE dilution. Individual symbols represent data derived from HC (n=6), ERAPB (n=6) or LRASF (n=6) (Treg/Tresp ratio 1:10). Results were comparable when using either of these methods. The potency of suppression was calculated as [1- (proliferation of Treg+Tresp coculture / proliferation of Tresp cells)]. *p<0.01 vs HC; † p<0.01 vs ERA, Mann Whitney test.

### Suppressive action of Treg cells on Tresp cell soluble TNFα, IFNγ and IL-17 secretion

Total Treg, isolated Treg39+ and isolated Treg39- cells were able to downregulate the amount of soluble TNFα and IFNγ secretion detected in coculture supernatants (Fig. 5A), and again the suppressive potency of ERA Total Treg cells was superior to that of HC Tregs (Fig. 5A), whereas LRASF Tregs demonstrated the highest potency (Fig. 5A). In addition, and paralleling the results observed in assays of proliferation, Treg39+cells were significantly more potent as compared with Treg39-, and the potency of Treg39+ or Treg39- was comparable among the groups (Fig. 5A), suggesting that the differences observed in assays using total Tregs can be attributable to the increased Treg39+ proportions present in ERA blood and LRASF. In contrast, the concentration of soluble IL-17 was not downmodulated in cocultures of Tresp with total Treg, Treg39+ or Treg39- cells (Fig. 5A) which is likely attributable to the fact that Tregs produce IL-17 themselves (Fig. 2 A, B; Fig. 3 A, B).

**Figure 5.**
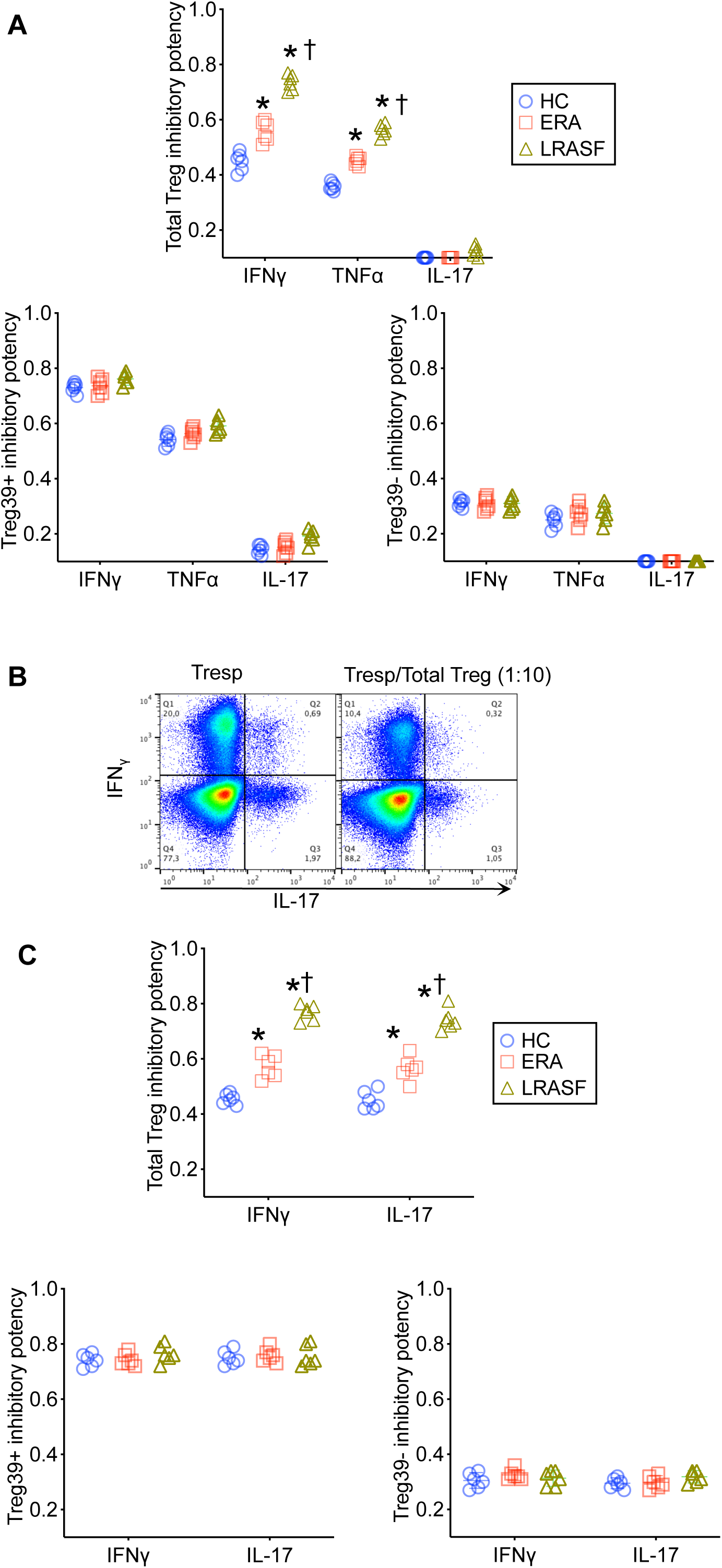
Inhibitory potency of Treg cells on Tresp cell cytokine secretion. The potency of inhibition of Treg on Tresp cell cytokine secretion was assessed in 6 day Treg/Tresp cell cocultures (Treg/Tresp cell ratio 1:10). Soluble cytokines were assayed by ELISA in culture supernatants and in addition, intracellular cytokine expression was examined by cytometry after establishing a gate in CFSE positive cells; this was possible because only Tresp cells were labeled with CFSE before the initiation of the cocultures. The potency of suppression was calculated as [1- (proliferation of Treg+Tresp coculture / proliferation of Tresp cells)]. **A. Inhibition of soluble cytokine concentrations in Treg/Tresp coculture supernatants.** Total Treg, isolated Treg39+ and isolated Treg39- cells were able to downregulate the concentration of soluble TNFα and IFNγ, but not of IL-17, detected in coculture supernatants. Individual symbols represent data derived from HC (n=6), ERAPB (n=6) or LRASF (n=6) (Treg/Tresp ratio 1:10). B. Inhibition of Tresp intracellular cytokine expression in Treg/Tresp coculture supernatants. Representative dot plots show IL-17 and interferon gamma expression in cells gated for CFSE. Total Treg, isolated Treg39+ and isolated Treg39- cells were able to downregulate the concentration of intracellular IFNγ and IL-17. Individual symbols represent data derived from HC (n=6), ERAPB (n=6) or LRASF (n=6) (Treg/Tresp ratio 1:10). *p<0.01 vs HC; † p<0.01 vs ERA, Mann Whitney test.

### Suppressive action of Treg cells on Tresp cell intracellular IFNγ and IL-17 secretion

Tresp cell intracellular secretion of IFNγ and IL-17, and their variation in the presence of Treg cells, were evaluated by cytometry after establishing a gate in CFSE positive cells (Fig. 5B); this was possible because only Tresp cells were labeled with CFSE before the initiation of the cocultures. In this setting, it was evident that Total Treg, isolated Treg39+ and isolated Treg39- cells were able to downregulate not only the amount of intracellular IFNγ but also of IL-17 secretion (Fig. 5 B, C). Once more, the suppressive potency of ERA Total Treg cells was superior to that of HC (Fig. 5 C), whereas LRASF Tregs demonstrated the highest potency (Fig. 5C). Also, Treg39+ were more potent than Treg39- cells, and the potency of Treg39+ or Treg39- was comparable among the groups (Fig. 5C), suggesting that the differences observed in assays using total Tregs can be attributable to the increased Treg39+ proportions present in ERA blood and LRASF.

### Effect of ADO pathway blockade on CD39+ and CD39- Treg cell potency

We then tested the effect of ADO pathway inhibitors in cocultures of total, CD39+ or CD39- Treg cells with CD39-Tresp cells (1:10 Treg/Tresp ratio), using soluble IFNγ secretion as a readout. These experiments were performed with cells derived from the peripheral blood of 6 HC and 6 treatment-naïve ERA patients. As stated above, total ERA Tregs were significantly more potent suppressors when compared with HC (Fig. 6). In the presence of the ADO receptor 2A (ADORA2A) antagonist ZM241385 or the CD39 inhibitor ARL67156 the potency of total Tregs was significantly reduced to a final level that was not different for ERA vs HC (Fig. 6); in contrast, the ADO receptor A1 (ADORA1) antagonist DPCPX did not modify these potencies (Fig. 6). The suppressor activity of Treg39+ cells was likewise abrogated by ZM241385 or ARL67156 but not by DPCPX (Fig. 6), to a similar extent in ERA and HC, whereas the function of Treg39- cells was not modified by any of these agents (Fig. 6).

**Figure 6.**
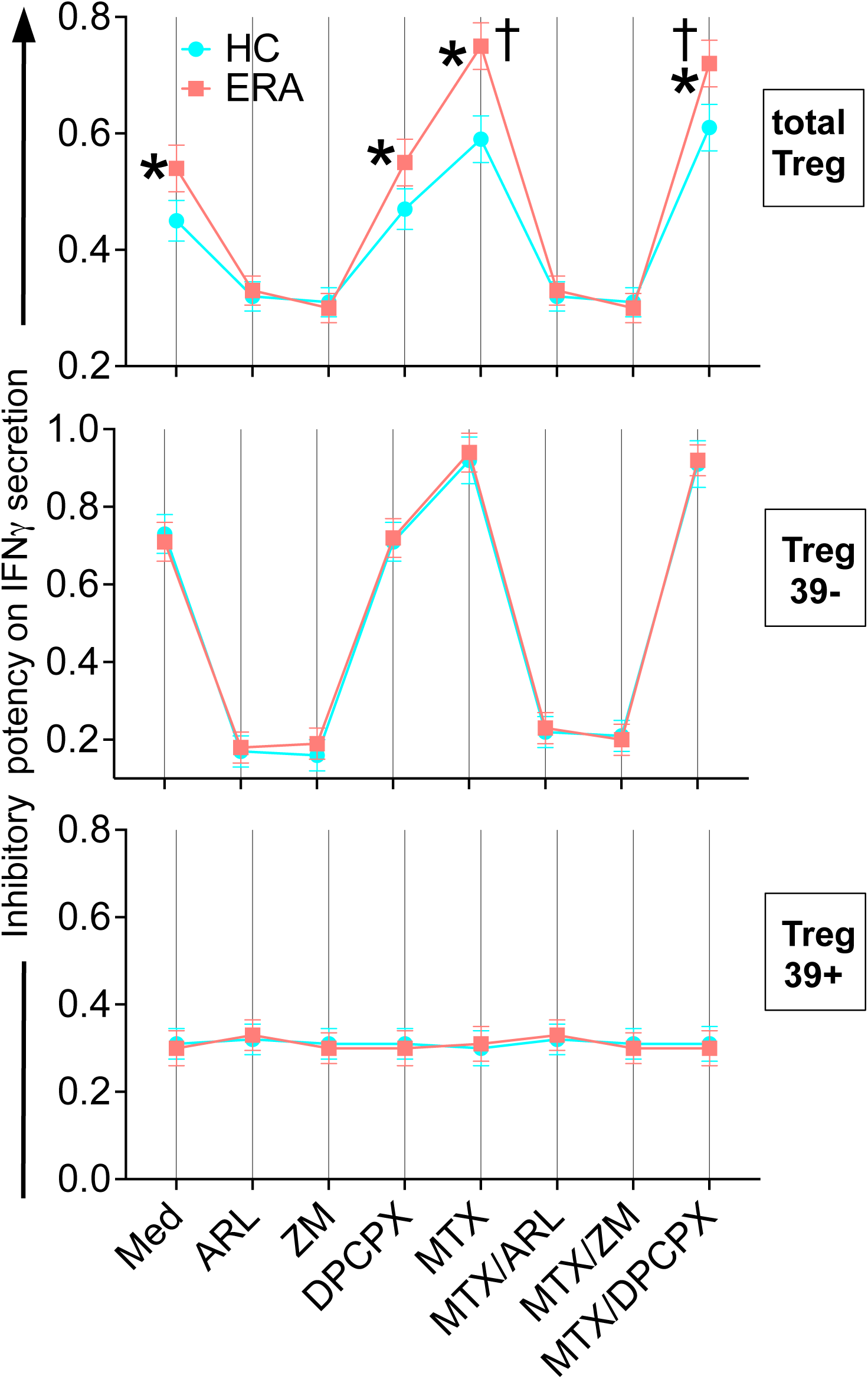
Effect of the CD39 inhibitor ARL67156, the ADORA2AR antagonist ZM241385 or the ADORA1 antagonist DPCPX on the potency of total Treg, CD39+ or CD39- Treg, in the presence or absence of MTX. *p<0.01 vs HC; † p<0.01 vs cells cultured in medium without MTX (Mann Whitney test).

### Effect of MTX on CD39+ and CD39- Treg cell function

In the presence of MTX, the suppressor potency of total Tregs or Treg39+ cells was significantly enhanced (Fig. 6). This enhancement was more marked for ERA total Treg as compared with HC Treg (Fig. 6) but similar for ERA Treg39+ as compared with HC Treg39+ cells (Fig. 6). No effect of MTX could be observed in the presence of ZM241385 or ARL67156, both of which significantly reduced the potency of total Treg or Treg39+ to levels that were comparable for both subpopulations; in contrast, DPCPX did not modify the action of MTX (Fig. 6). In addition, MTX did not modify the suppressor potency of Treg39- cells. This again suggests that the differences observed in assays using total Tregs can be attributable to the increased Treg39+ proportions present in ERA.

### Relation of the baseline circulating Treg and Treg39+ cell frequencies in ERA with response to treatment

A total of 63 ERA patients were re-evaluated clinically 12 months after initiating treatment with MTX and low-dose prednisone. At that time point, 39 patients (62 %) demonstrated low disease activity (LDA) as determined by a DAS28-ESR ≤3.2 [28,29]; all of them had also experienced a drop of their basal DAS28 (ΔDAS28) >1.2, hence had achieved a good EULAR response [29]. The remaining 24 patients (38 %) showed a moderate disease activity (3.2< DAS28-ESR ≤ 5.1) and moderate EULAR response (ΔDAS28 >1.2) [29]. Patients who achieved LDA had demonstrated significantly higher basal frequencies of both total Treg [OR 2.65 (95% CI, 1.65-4.9)] and Treg39+ cells [OR 1.93 (1.33-4.5)] as compared to the group with moderate activity (Fig.7A), and multiple logistic regression indicated that this was independent of age, basal disease activity, RF or ACPA titres. The Receiver Operating Characteristic (ROC) analysis demonstrated an area under the curve (AUC) of 0.85 (95% CI, 0.76-0.95) for the association of total Treg with response and 0.98 (0.94-1.0) for Treg39+ cells (Fig. 7A). The relative risk (RR) of achieving LDA for patients with a circulating Treg cell frequency > 7.6 (p75 value observed in HCs) was 3.2 (95% CI 1.9-6.0) and of patients with a circulating Treg39+ frequency > 39.3 (75th percentile of the values observed in HC) was 13.4 (2.9-75.6) (Fisher’s exact test).

**Figure 7.**
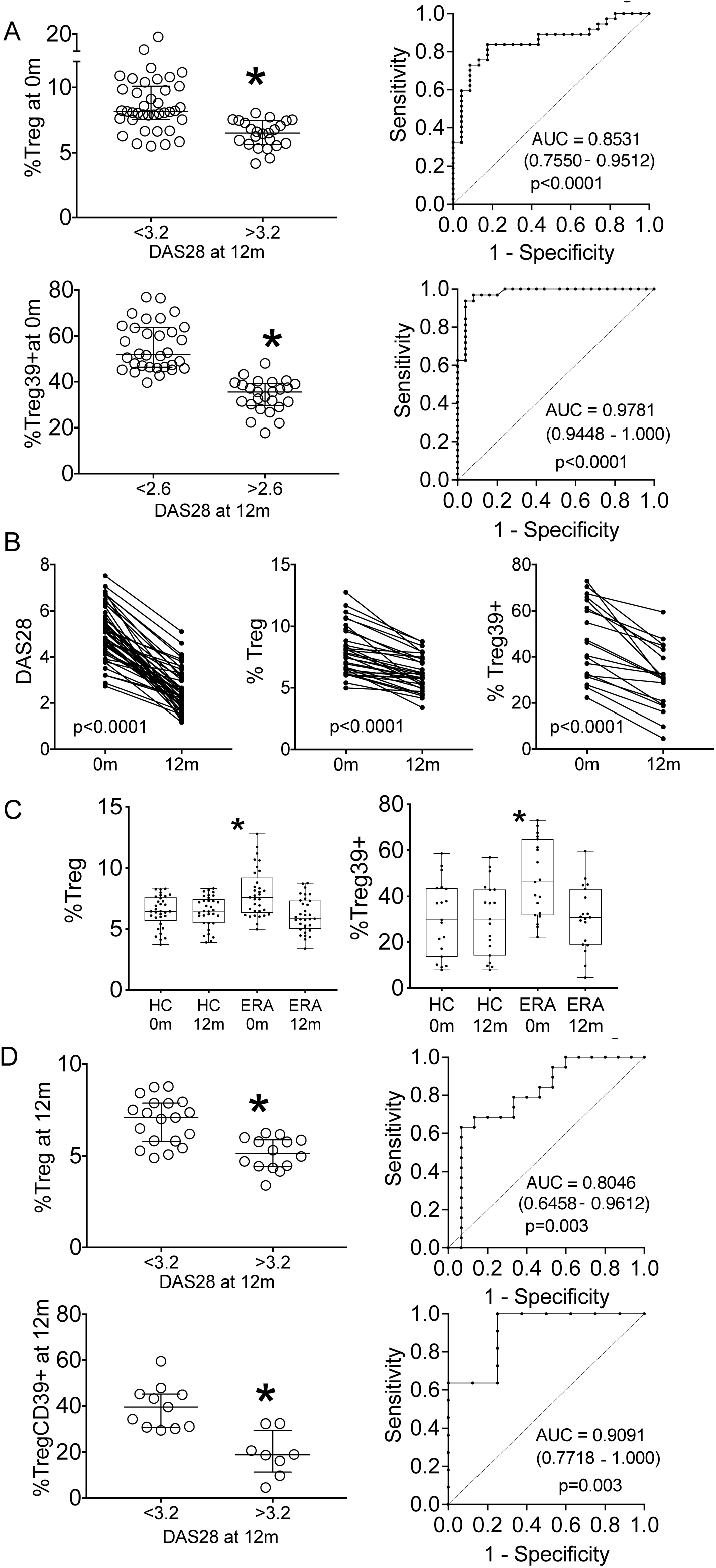
Relation of the clinical response with the basal or 12-month circulating total Treg or Treg39+ cell frequencies in ERA. **A**. Pretreatment frequency of circulating total Treg or Treg39+ cells in 63 ERA patients who did did (n=39) or did not (n=24) achieve low disease activity (LDA) 12 months after inititating methotrexate (LDA defined as DAS28≤ 3.2) (Bars represent the median and interquartile range; *p<0.05, Mann Whitney test) and associated ROC curves. **B**. DAS 28 (n=63), and frequency of total Treg (n=34) or Treg39+ cells (n=19) at the basal and 12 month visit in ERA patients (p values calculated using the Wilcoxon signed rank test). **C**. Frequency of Total Treg (n=34 for HC and ERA patients) or CD39+ Treg cells (n=19) for HC and ERA patients), at the basal and 12 month visit; box and whiskers represent the median, interquartile range, maximum and minimum values; *p<0.05 vs HC (Mann-Whitney test). **D**. Frequency of total Treg (n=34) or CD39+ Treg (n=19) at the 12 month visit, in patients who had (Treg, n=19; Treg39+, n=11) or had not (Treg, n=15; Treg39+, n=8) achieved low disease activity (LDA) (Bars represent the median and interquartile range; *p<0.05 vs HC, Mann-Whitney test), and associated ROC curves.

### Modifications of the circulating Treg and circulating Treg39+ cell frequency in ERA along the disease course

The frequency of circulating Tregs was re-evaluated 12 months after the first visit in 34 ERA patients [19 of whom (56%) demonstrated LDA], and the frequency of Treg39+ cells was re-examined in 19 of these subjects [11 of whom (58%) showed LDA]. At the 12 month visit, all of the patients were receiving oral MTX at doses of 15–25 mg/week. Prednisone had been discontinued in 25 subjects and the remaining 8 patients, all of whom demonstrated moderated disease activity, were taking 2.5-5 mg of prednisone daily. A significant decrease of the basal DAS28 (Fig. 7 B) was accompanied by a significant reduction of the initially elevated circulating Treg and Treg39+ cell frequencies in ERA patients (Fig. 7 B) with no variation observed in HC (Fig. 7 C), and differences between HC and ERA were no longer observed (Fig. 7 C).

### Relation of the 12-month circulating Treg and circulating Treg39+ cell frequency in ERA with response to treatment

Interestingly, despite the observed drop in their numbers, the circulating Treg [OR 2.3 (1.3-4.9)] and Treg39+ [OR 1.4 (1.1-2.7)] cell frequencies observed at the 12 month visit were also associated with the EULAR response (Fig. 7 D). Because, as opposed to the basal study, at the 12 month examination patients were receiving MTX with or without low-dose prednisone, it is of interest to determine whether these drugs are conditioning the observed Treg or Treg cell frequencies at 12 months. Of note, since none of the 19 patients with LDA were taking steroids at this time point and 8 out of the 11 patients (73%) with moderate disease activity were taking them, the intake of steroids in this group of patients acts as a confounder that cannot be discerned from disease activity a in multivariate analysis aimed at estimating the relation of disease activity with Treg cell frequencies and therefore we are unable to determine whether prednisone is modifying the Treg or Treg39+ cell frequencies. In addition, there were no differences in the dose of MTX that was being administered to the patients who achieved LDA as compared with those who did not; and therefore we could not conclude whether the dose of MTX was associated with the Treg or Treg39+ cell frequencies at 12 months.

### Effect of RA Synovial Fibroblasts on Treg CD39 and Treg IL-17 expression

We previously described that RASFib constitutively express surface IL-15 and are able to augment the potency of isolated Treg cells [6]; hence we next tested their effect on Treg CD39 and IL-17 expression. As shown in Fig. 8, isolated Treg cells significantly upregulated their CD39 and downregulated their IL-17 expression when cocultured with RASFib in the presence of an anti-CD3ε MoAb; this effect required direct cell contact since it was not observed in the presence of transwell inserts (Fig. 8), and was abrogated by a neutralizing anti IL-15 antibody (Fig. 8) but not by the binding control antibody W6/32 (anti HLA class I) (Fig. 8) or the additional negative controls anti-MHC class II or isotype control murine IgG1 (not shown). This modulation of CD39 and IL-17 expression was not observed with anti-CD3ε MoAb alone (Fig. 8) or in cocultures with DFib or OASFib (Fig. 8) that do not constitutively express surface IL-15 [6].

**Figure 8.**
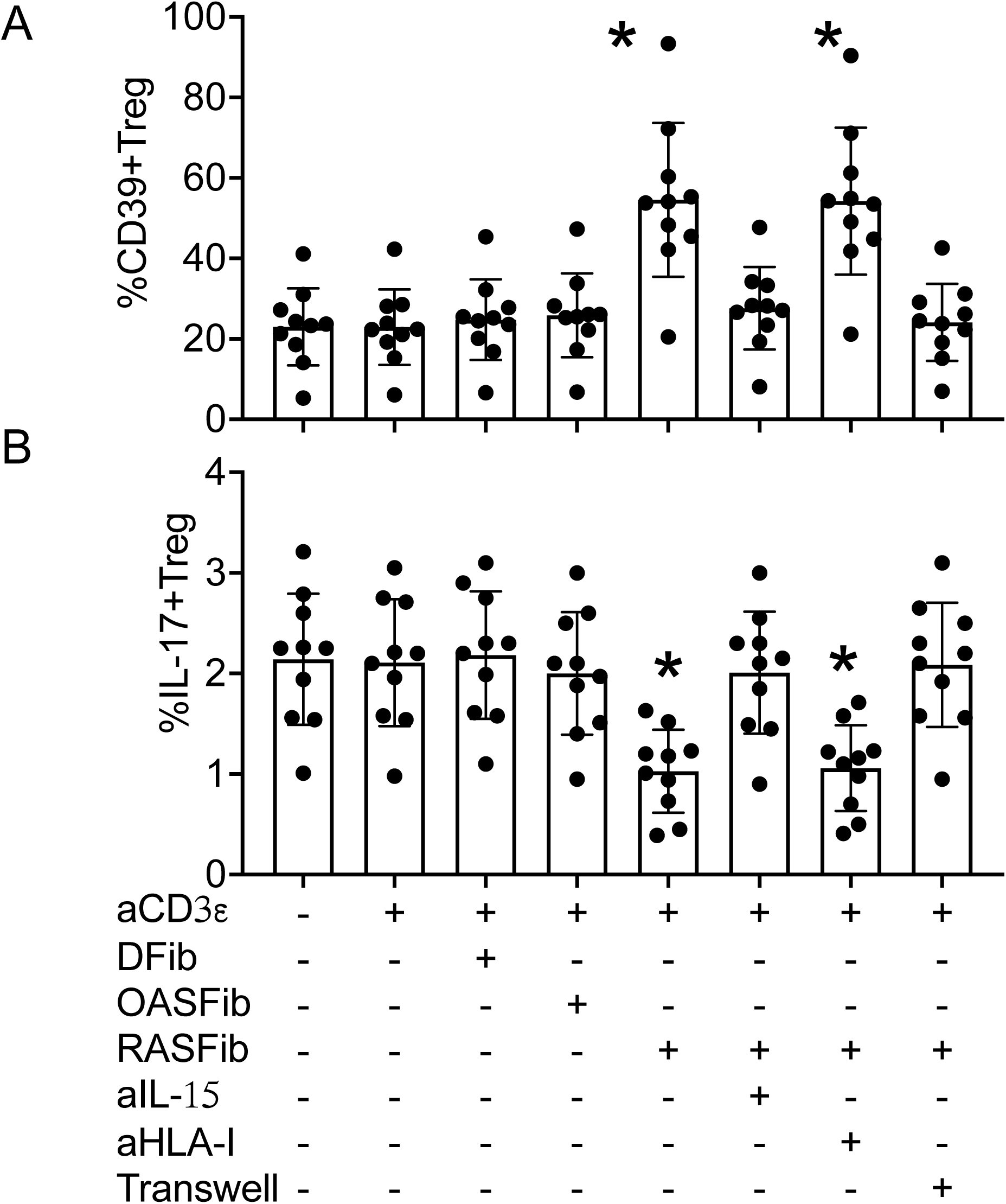
Modulation of Treg CD39 and IL-17 expression in coculture with RA synovial fibroblasts (RASFib). Percentage of CD39+ (A) or IL-17+ (B) expressing cells among total regulatory T cells isolated from 10 healthy controls, right after isolation (far left columns), or after culturing for 5 days in medium supplemented with an aCD3ε mAb, in the presence or absence of human dermal fibroblasts (DFib), osteoarthritis synovial fibroblasts (OASFib) or RA synovial fibroblasts (RASFib). In some conditions, cocultures of Tregs with RASFib were supplemented with a neutralizing anti-IL-15 mAb, or a binding control anti-HLA class I antibody, or performed using transwell inserts that prevent direct contact between fibroblasts and Treg cells. *p<0.05 vs expression of CD39 or IL-17 in freshly isolated Tregs (Mann-Whitney Test).

## DISCUSSION

Regulatory T cells (Tregs) play a key role in the control of inflammatory conditions [1–12] and a better understanding of their pathophysiology may help design new personalized therapeutic strategies. Methotrexate (MTX) remains the drug of choice for the initial management of Rheumatoid Arthritis (RA) [16] but around 30% of the patients discontinue treatment due to an insufficient response or adverse events [16,30], and there is a pressing need for reliable biomarkers that would help identify good responders.

Despite intensive research there is still controversy regarding the frequency, phenotype and function of regulatory T (Treg) cells in established [7–9,31,32] or early RA [6,11,12,33,34]. Frequencies have been shown to be, when compared with HC, either increased [6,9,33,34], akin [31] or decreased [11,12]. In addition, Treg cell function has been described as increased in patients with ERA [6] and in mice at the early phase of experimental RA [35] but also decreased [7,32] or similar [11,31] to HC in patients with early or longstanding RA. Of note, Lawson et al [11] reported decreased proportions of circulating Tregs in ERA with preserved functional capacity; in contrast, Cribbs et al found that the function but not the number of Treg cells was altered in DMARD-naïve RA [7]; other investigators have reported decreased [12] or increased [33,34] numbers of circulating Tregs in ERA but did not study their suppressive capacity. In addition, Cribbs et al. [7] described that treatment with DMARDs normalized the altered Treg cell function in previously untreated patients and at the same time induced an elevation of Treg cell numbers above those observed in HC [7]; however, Ehrenstein et al. showed that whereas DMARDs were not able to restore the altered Treg cell function or to elevate Treg cell numbers in RA above HC levels, anti-TNF drugs did achieve both of these objectives [32]. Differences among studies may be related to discordant characteristics of the subjects given the clinical heterogeneity of RA, or alternatively to technical issues such as cytometry methods or experimental settings of the functional studies.

We have herein examined the features of Tregs in patients with untreated early RA (ERA) thereby eliminating the possible interference of immuno-modulatory drugs or chronic inflammatory feedback mechanisms, and compared them with circulating and synovial fluid Tregs of patients with longstanding RA.

We previously reported that ERA patients demonstrate an increased frequency of circulating Treg cells [6] with an enhanced suppressor activity [6], which is confirmed and extended by the data presented herein: ERA but not LRA Tregs show increased proportions of Treg39+ together with decreased proportions of Treg/Th17 cells, and functional studies indicate that this redistribution of subsets seems to account for the superior ERA Treg cell potency. Interestingly, the Treg cell subset composition differs between early and more advanced stages of the disease. At the same time, CD39 was maximally increased and IL-17 expression further decreased in RA synovial fluid Tregs, suggesting that the inflammatory microenvironment may modulate the phenotype of the local Tregs.

Tregs are potent suppressors of responder T cell function [1, 2]. However, previous investigations regarding the capacity of Treg39+ cells to inhibit IL-17 secretion have yielded conflicting results [5, 21–23]. We observed that Treg39+ and Treg/Th17 are not distinct but overlapping subsets: IL-17 secretion was detected by cytometry in both Treg39+ and Treg39- cells, although Treg39+ contained an inferior proportion of IL- 17+ cells as compared with Treg39- cells of the same group of subjects. Interestingly, when compared with HC Treg39+, ERA Treg39+ cells contained a lower percentage of IL-17+ cells whereas RASF had the lowest IL-17 expression (Fig. 2B); in contrast, IL- 17+ cells were equally represented among HC, ERA or LRA Treg39- cells. This indicates that the reduced IL-17 secretion of ERA or RASF total Tregs is not only related to their higher content of Treg39+ but also to the lower IL-17 production by their Treg39+ cells. Since Tregs produce IL-17 themselves, the concentration of soluble IL-17 detected in supernatants was not downmodulated in cocultures of Tresp cells with total, CD39+ or CD39- Tregs. However, studies using flow cytometry of Tresp cells stained with CFSE revealed that all three Treg cell populations were able to inhibit Tresp cell intracellular IL-17 secretion; this adds to the undestanding of Treg cell biology; nonetheless, from a pathophysiological point of view, the conclusion that IL- 17 levels in the inflammatroy milieu cannot be controlled by Treg cells remains. In addition, all three Treg cell populations were potent suppressors of Tresp proliferation, TNFα and IFNγ secretion. Indeed, functional assays revealed that, as compared with HC, total ERA Treg cells displayed an enhanced suppressor capacity while the potency of RASF Tregs was maximal, consistent with our previous observation [6]. Of note, the suppressor potency of isolated Treg39+ or Treg39- cells was not significantly different among the studied groups, whereas Treg39+ were much more potent suppressors than Treg39- cells. This suggests that the observed superior strength of total ERA or RASF Tregs can be attributable to their higher relative amounts of Treg39+ cells.

The mechanisms mediating the suppressor function of Treg cells are not well understood and CD39 expression seems to play an important role [14,15,17]. Indeed, we observed that ADORA2A or CD39 inhibitors significantly reduced the action of total or Treg39+ cells but had no effect on Treg39- cells. Additionally, in the presence of MTX the suppressor potency of total or Treg39+ was further increased and this enhancement was hindered by ADORA2A or CD39 blockade; in contrast, MTX did not significantly modify the suppressor capacity of Treg39- cells. This indicates that, consistent with previous reports [8,14,15], Treg39+ cells exert their suppressor effect through the enzymatic activity of CD39, and also implies that MTX may cooperate with Treg39+ by providing an additional source of extracellular ADO precursors via ATIC inhibition [17,18]. In this context, whereas neutralization of CD39 activity or ADORA2A signaling have proved effective against cancer [36], sinergizing with this mechanism is a potential therapeutic strategy in autoimmunity, and new drugs based in this mode of action could be beneficial for RA patients who develop unwanted side effects of MTX.

Remarkably, the elevated basal proportions of both total Treg and Treg39+ cells were associated with the clinical response to methotrexate at 12 months, and the RR of achieving LDA (DAS28-ESR≤3.2) for patients with a basal Treg39+ frequency >39.3 (p75 seen in HC) was 13.4 (95% CI, 2.9-75.6), which is consistent with data reported by other groups in LRA [8,19] or ERA [20]. This suggests that, in parallel with the observed in vitro results, MTX can also cooperate with Treg39+ cells in vivo.

We herein report for the first time that not only were total peripheral blood Treg and Treg39+ cell frequencies augmented in ERA but interestingly, 12 months after initiating treatment with MTX, both frequencies had significantly dropped and were no longer above HC levels. Furthermore, their association with the clinical response remained: indeed, the circulating Treg and Treg39+ cell frequencies observed at 12 months were still significantly higher in those patients who had achieved LDA. This suggests that not only Treg cell numbers [6] but also Treg cell expression of CD39, are upregulated during the initial stages of ERA as a negative feedback mechanism aimed at controlling inflammation. Of note, the expression level of CD39 on Tregs is genetically determined [37] but can be overinduced if exposed to an inflammatory milieu [37]. In this context, the transient expansion of Treg39+ cells in ERA may provide a therapeutic window during which methotrexate has a superior effectivenes.

Indeed, in a well characterized murine model of experimental arthritis, K. Monte et al [35] showed an expansion of Treg cell numbers and function at the early phases, with dynamic changes along the course of the disease [35] which parallels the results we are showing herein. In this context the inflamed joint microenvironment, by interacting with infiltrating Tregs, can play an important role in modulating their phenotype, and function.

RA synovial fibroblasts display a distinctive phenotype that confers them the capacity to erode bone and cartilage [38,39]. In addition, they establish direct interactions with lymphocytes thereby contributing not only to initiate and perpetuate [38–41] but also to regulate the chronic RA inflammation [6]. Specifically RASFib, like human spleen myofibroblasts [42] and bone marrow stromal cells [43], constitutively present IL-15 on their cell membrane [6,40,41] that has a dual action on the equilibrium between regulatory and effector T cells [6]. We herein report that when isolated Tregs from the peripheral blood of HC were cocultured with RASFib, not only did they augment their suppressive capacity as previously described by our group [6]: at the same time their CD39 expression was upregulated and in parallel their IL-17 expression was downregulated thereby acquiring a phenotype closer to RA synovial fluid Tregs. Neutralization experiments indicated that this effect is facilitated by RASFib surface IL-15 expression and is in line with publications indicating that IL-15 supports Treg cell stability [6,44,45] while simultaneously curtailing Th17 cell generation [44,46], and with observations manifesting that the phenotype of RA synovial fluid T cells can be modulated by their interaction with synovial fibroblasts in the lining layer [47]. Accordingly, the number and phenotype of peripheral blood Tregs we observed in ERA may reflect Treg recirculation from and to inflamed sites [48,49].

In summary, MTX enhances the potency of Treg39+ cells in vitro, and the pretreatment frequency of circulating total Treg39+ cells can help identify good responders to MTX thereby contributing to the development of personalized medicine strategies. In addition, the expanded circulating Treg39+ cell proportions in untreated early ERA may provide a window of opportunity facilitating the effectiveness of prompt MTX initiation.

## Patient and Public Involvement statement

Patients or the public WERE NOT involved in the design, or conduct, or reporting, or dissemination plans of our research

## AUTHOR CONTRIBUTIONS

All authors were involved in drafting the article or revising it critically for important intellectual content, and all authors approved the final version to be published. Dr. Miranda-Carús had full access to all of the data in the study and takes responsibility for the integrity of the data and the accuracy of the data analysis.

**Study conception and design.** Miranda-Carús

**Acquisition of data**. Villalba, Nuño, Benito-Miguel, Novella-Navarro, Monjo, Peiteado, García-Carazo, Balsa, Miranda-Carús

**Analysis and interpretation of data**. Miranda-Carús, Benito-Miguel, Nuño, Villalba, Monjo, Novella-Navarro, Peiteado, García-Carazo, Balsa

**Wrote the manuscript**. Miranda-Carús

## DATA AVAILABILITY STATEMENT

The raw data supporting the conclusions of this article will be made available by the corresponding author upon reasonable request.

## KEY MESSAGES

**What is already known about this subject?**

- Treg cells are essential for restraining autoimmunity
- Provision of adenosine by CD39+ T reg cells (Treg39+) significantly contributes to suppression
- Methotrexate (MTX), via ATIC inhibition, provides adenosine precursors to be metabolized by CD39

**What does this study add?**

- In vitro, MTX augments the potency of Treg39+ cells
- The pretreatment circulating Treg39+ frequency in ERA can help identify good responders to MTX
- Early RA (ERA) patients demonstrate transiently increased Treg39+ cells which can provide a slot for prompt treatment initiation

**How might this impact on clinical practice or future developments?**

The circulating Treg 39+ cell frequency is a potential tool for personalized medicine strategies in ERA.

